# Pleiotropic function of the *oca2* gene underlies the evolution of sleep loss and albinism in cavefish

**DOI:** 10.1101/2020.09.27.314278

**Authors:** Morgan O’Gorman, Sunishka Thakur, Gillian Imrie, Rachel L. Moran, Erik Duboue, Nicolas Rohner, Suzanne E. McGaugh, Alex C. Keene, Johanna E. Kowalko

**Affiliations:** Jupiter Life Science Initiative, Florida Atlantic University, Jupiter, FL 33458; Department of Ecology, Evolution, and Behavior. University of Minnesota, St. Paul, MN 55108; Stowers Institute, Kansas City; Department of Biology Science, Florida Atlantic University, Jupiter, FL 33458; Harriet L. Wilkes Honors College, Florida Atlantic University, Jupiter, FL 33458

## Abstract

Adaptation to novel environments often involves the evolution of multiple morphological, physiological and behavioral traits. One striking example of multi-trait evolution is the suite of traits that has evolved repeatedly in cave animals, including regression of eyes, loss of pigmentation, and enhancement of non-visual sensory systems [1,3]. The Mexican tetra, *Astyanax mexicanus*, consists of fish that inhabit at least 30 caves in Northeast Mexico and ancestral-like surface fish which inhabit the rivers of Mexico and Southern Texas [6]. Cave *A. mexicanus* are interfertile with surface fish and have evolved a number of traits that are common to cave animals throughout the world, including albinism, eye loss, and alterations to behavior [8–10]. To define relationships between different cave-evolved traits, we phenotyped 208 surface-cave F2 hybrid fish for numerous morphological and behavioral traits. We found significant differences in sleep between pigmented and albino hybrid fish, raising the possibility that these traits share a genetic basis. In cavefish and many other species, mutations in *oculocutaneous albinism 2* (*oca2*) cause albinism [11–15]. Surface fish with CRISPR-induced mutations in *oca2* displayed both albinism and reduced sleep. Further, this mutation in *oca2* fails to complement sleep loss when surface fish harboring this engineered mutation are crossed to different, independently evolved populations of albino cavefish with naturally occurring mutations in *oca2*, confirming that *oca2* contributes to sleep loss. Finally, analysis of the *oca2* locus in wild caught cave and surface fish suggests that *oca2* is under positive selection in at least three cave populations. Taken together, these findings identify *oca2* as a novel regulator of sleep and suggest that a pleiotropic function of *oca2* underlies the adaptive evolution of both of albinism and sleep loss.

## Results

Colonization of a novel environment often results in evolution of behavioral, morphological, and physiological traits. Uncovering the relationships between these traits is critical to understanding the mechanisms that drive adaptation. Multiple populations of cave *A. mexicanus* have evolved a suite of traits that distinguish them from surface dwelling counterparts. These include regression of eyes, albinism, elevated energy stores, and sleep loss [4,16–19]. To investigate the relationship between evolved traits in cave *A. mexicanus*, we generated hybrids from surface fish and cavefish from the Pachón cave, which are highly troglomorphic. Surface-Pachón F2 hybrids possess variable morphological phenotypes (Fig 1A). We quantified numerous traits associated with cave evolution including eye size, albinism, adiposity, feeding and sleep [4,8,17–19]. At 20 dpf, surface fish, cavefish, and surface-cave F1 and F2 hybrids were assayed for sleep over a 24hr period, followed by measurements of food consumption. Fish were then stained with Nile Red, which labels adipocytes in larval fish [19,20] and imaged to score pigmentation, adiposity, and eye diameter (Fig 1B). As individual fish were followed throughout all phenotyping steps, this experimental design allows for determining the relationship between traits in F2 hybrid fish.

**Figure 1.**
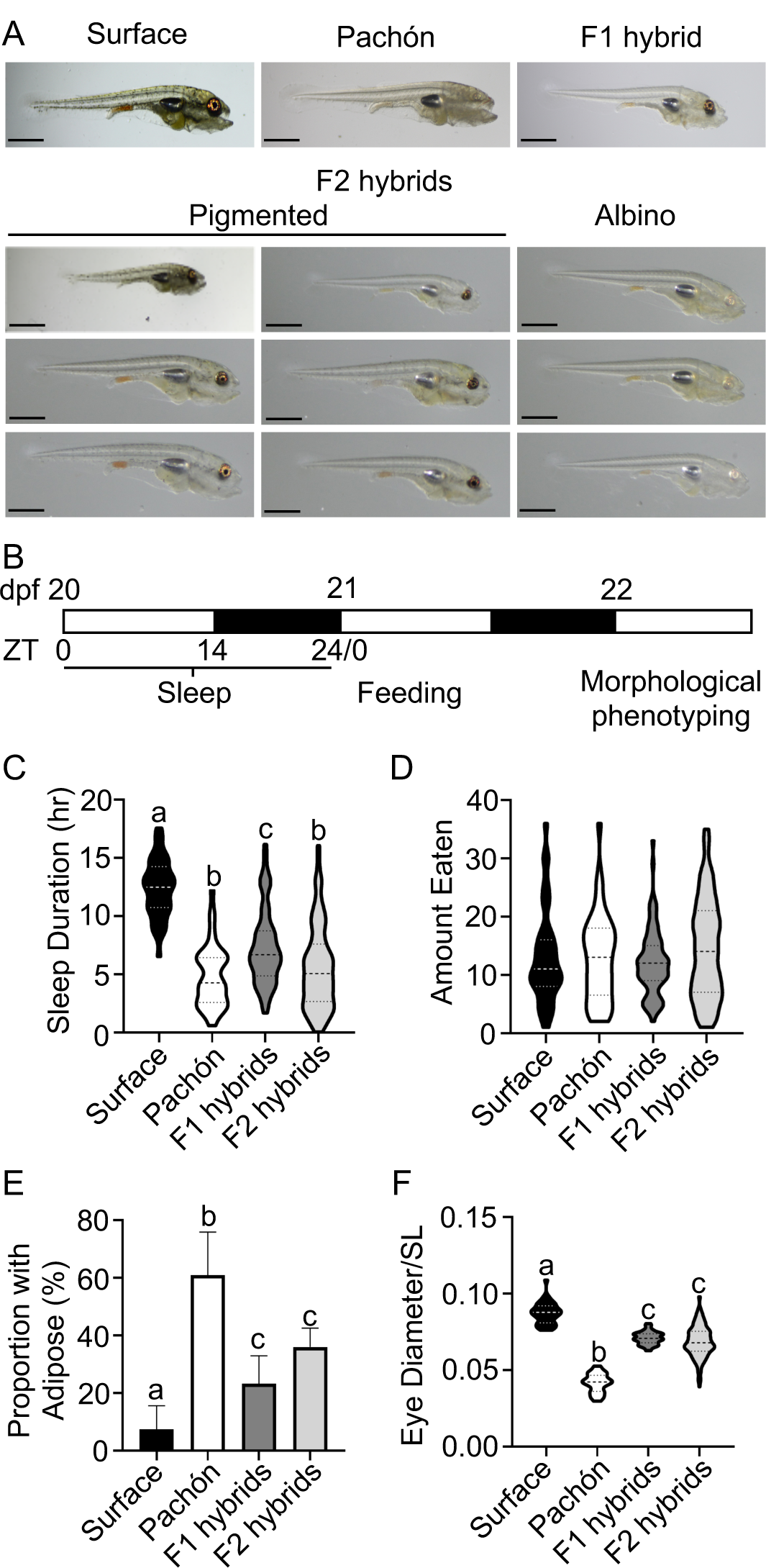
Genetic analysis of multiple traits evolved in cave *A. mexicanus*. (A) Images of 22dpf fish (scale = 1mm). (B) Timeline for phenotypic analysis. Surface fish (n=40), Pachón cavefish (n=41), surface-Pachón F1 hybrids (n=73) and surface-Pachon F2 hybrid fish (n=208) were phenotyped for (C) Total sleep duration (Kruskal-Wallis: H_2_=97.37, p <0.0001; Dunn’s multiple comparison: Pa vs. F1: z=3.55, p= 0.002, F1 vs. F2: z=3.95, p=0.0016). All other: p<0.0001) (D) Total brine consumed (Kruskal-Wallis: H_2_=4.36, p=0.2256 (E) Proportion of individuals with adipose. Error bars calculated using z*-value of 1.96 and denote the margin of error of the sample proportion. Fishers Exact tests: SF v Pa p<0.0001, SF v F1 p=0.0408, SF v F2 p=0.0002, Pa v F1 p=0.0001, Pa v F2 p=0.0048, F1 v F2 p=0.0590). (F) Eye diameter, corrected for standard length (ANOVA: F=197.4, p<0.0001; Tukey’s: F1 vs. F2 q=5.76, p=0.1970. all others p<0.0001). Statistical differences are indicated by different letters in (C)-(F).

Comparison of hybrids to pure cave and surface populations revealed variability suggestive of both monogenic and polygenic inheritance of the various traits, largely in agreement with previously published findings [4,21,22]. Sleep duration was significantly reduced in Pachón cavefish compared to surface fish. F1 hybrids slept more than pure Pachón cavefish and less than surface fish (Fig 1C, Fig S1A). F2 individuals were variable in the amount they slept, with some F2 fish sleeping very little, similar to Pachón cavefish, and other F2 fish sleeping at surface fish-like levels (Fig 1C). Additionally, sleep architecture was different between the groups. Sleep loss in Pachón cavefish is due to reductions in both the number and duration of sleep bouts relative to surface fish (Figure S1B,C). The average duration of each sleep bout in F1 hybrid fish was similar to cavefish, while bout length in F1 hybrids was intermediate between the surface fish and cavefish (Figure S1B,C). The dominant cave phenotype for bout length, but not bout number, raises the possibility that different components of sleep architecture are independently regulated. Indeed, bout number and bout duration were weakly correlated in F2 fish, (Spearman’s rho = 0.36, p<0.0001), consistent with at least some different genetic factors contributing to these different components of sleep architecture (Fig S1D).

One hypothesis for the evolution of sleep loss in cavefish is that altered foraging and/or metabolic demands associated with living in a food-poor environment drive the evolution of sleep loss [23]. In agreement with the previous findings that feeding does not differ between surface fish and Pachón cavefish [18,24], we observed no significant differences in total food consumption between surface fish, Pachón cavefish, F1 hybrids or F2 hybrids (Fig 1D). These results suggest that in the Pachón population, sleep loss is unlikely to be linked to changes in food consumption at the larval stage. The initial production of adipose deposits is ontogenically regulated and occurs earlier in Pachón cavefish relative to surface fish [19]. Within all four experimental groups, some fish had developed adipose deposits by 22 dpf, while others had not (Fig 1E, Fig S1E,F). However, a significantly larger percentage of cavefish had developed adipose deposits when compared to surface fish at 22 dpf. An intermediate percentage of F1 and F2 hybrid fish had developed adipose deposits at this stage relative to both cave and surface populations (Fig 1E), suggesting that differences in adiposity have a genetic basis and that regulation of adiposity shows partial or incomplete dominance.

Cave animals are characterized by loss of eyes and pigmentation [1]. Similar to previously reported studies [25,26], we found that Pachón cavefish eyes were significantly smaller than eyes in surface fish. Further, eyes in F1 hybrid fish were intermediate in size and significantly different from both parental populations (Fig 1F). Eye size ranged in F2 hybrid fish from surface-like to cave-like (Fig 1F). Therefore, nearly all traits analyzed with differences between surface fish and cavefish display intermediate phenotypes in F1 hybrids and variability in F2 fish consistent with a polygenic basis for evolved differences between *A. mexicanus* cave and surface populations.

Finally, we analyzed albinism, the complete loss of melanin pigmentation. All surface and F1 fish exhibited robust melanin pigmentation, while all cavefish were albino, consistent with the recessive nature of this trait [11,17] (Fig 1A). In our F2 population, 61 of the 208 fish tested were albino (Fig 1A). To examine segregation of albinism in F2 individuals precisely, we also quantified the number of albino F2 fish from a single clutch by collecting a random cohort of fish pre-pigmentation, and allowing these fish to develop until 5 days post fertilization, when albinism can be easily scored by eye as complete absence of melanin pigmentation. Of these fish, 66 out of 283 were albino (23.3%), which was not significantly different from the expected ratios for a trait controlled by a single gene (Chi square: p=0.69), consistent with the monogenic inheritance of albinism previously reported [11,17].

Cave traits could have evolved independently from each other, or through a shared genetic or functional basis. To determine if there is a relationship between cave-evolved traits, we performed pairwise comparisons between the traits that were significantly different between cavefish and surface fish: sleep, eye size, pigmentation and the presence of adipose tissue, reasoning that traits that were co-inherited would be significantly correlated with one another. We found three significant relationships from this analysis. F2 fish with adipose deposits had significantly smaller eyes compared to F2 fish without adipose (Cohen’s d=0.31, Table 1). Further, albino F2 fish had significantly smaller eyes than pigmented fish, raising the possibility that shared genetic architecture regulates these morphological traits (Cohen’s d=0.61, a moderate effect, Fig S2A, Table 1). Additionally, total sleep duration was significantly less in albino F2 fish compared to pigmented F2 fish (Cohen’s d=0.55, a moderate effect, Fig 2A,B). In albino F2 hybrid fish, both bout duration and bout number were significantly reduced, raising the possibility of shared genetic factors underlying the evolution of albinism and both sleep components (Fig 2C,D). These results suggest that albinism is associated with multiple traits that are present in cavefish populations. As little is known about the relationship between morphological and behavioral evolution, we focused on the relationship between albinism and sleep loss to determine if the co-heritability of these traits is due to closely linked loci or pleiotropy.

**Table 1.**
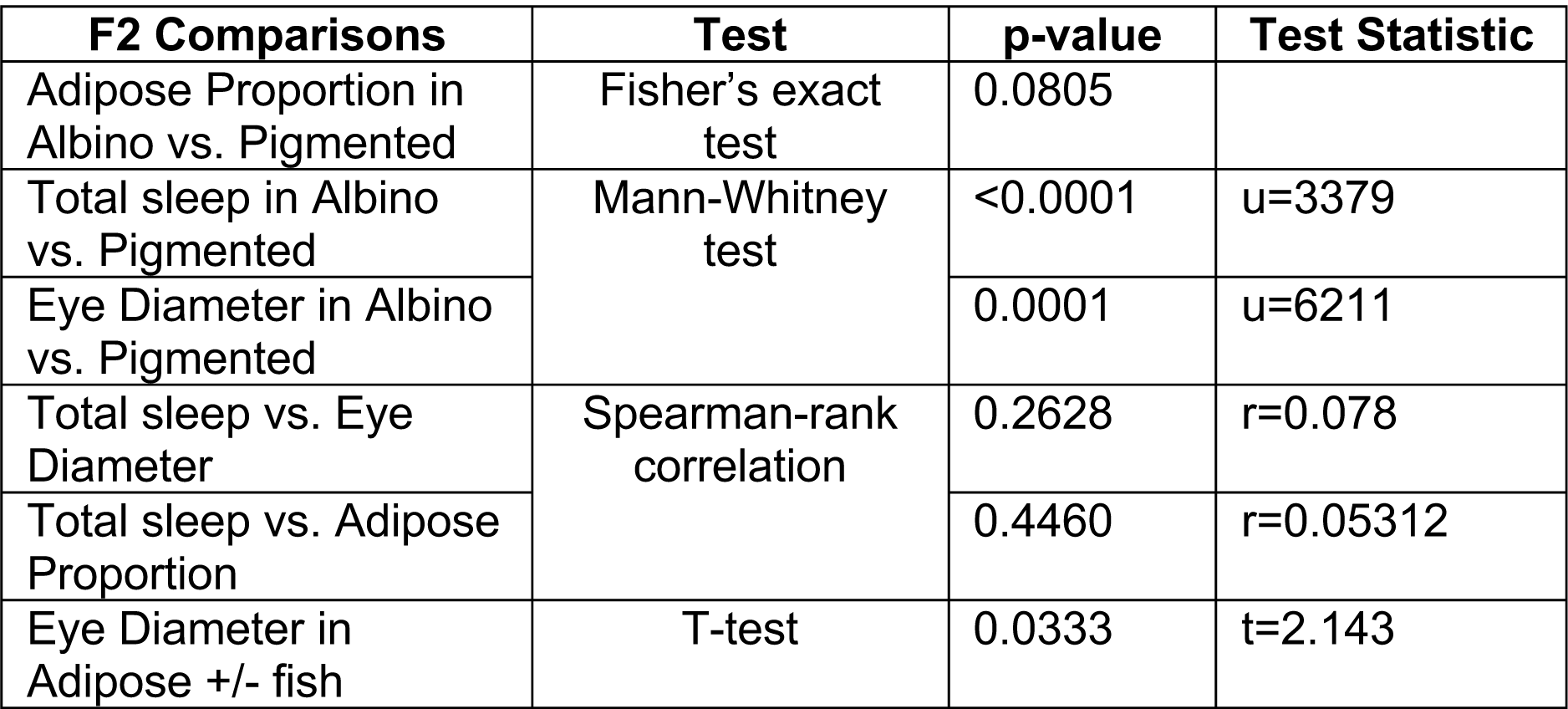
Pairwise comparisons of phenotypes in F2 individuals

**Figure 2.**
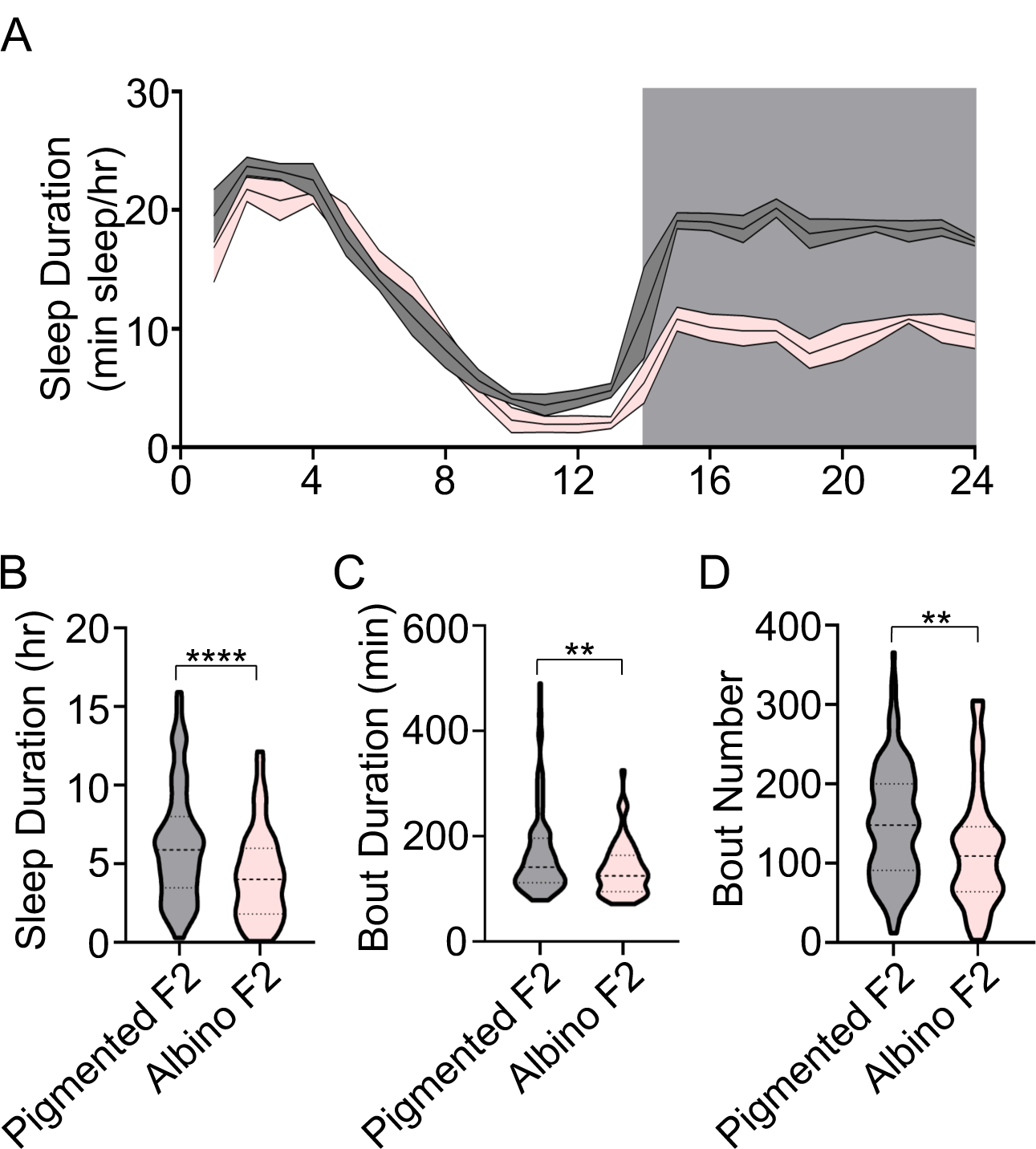
Relationship between sleep and pigmentation in cave-surface hybrid fish. (A) 24-hour sleep profile of F2 hybrid crosses depicting the amount slept at 1 hour intervals over a 24-hour time frame. The gray area of each graph represents laboratory night, when lights were off. The gray line represents pigmented F2s while the pink line represents albino F2s (B) Total sleep duration in albino compared to pigmented F2 hybrids (Mann-Whitney, U=3379, p<0.0001). (C) Bout duration of albino compared to pigmented F2 hybrids (t-test, t=2.67, p=0.0079). (D) Bout number in albino versus pigmented F2 hybrids (t-test, t=3.24, p=0.0043). For (A)-(D), pigmented F2 hybrids (n = 147), and albino F2 hybrids (n = 61). Graph (A) represents the average and standard deviation. Graphs (B)-(D) represent the median ± the quartile

In two independently evolved populations of cavefish, albinism is caused by deletions in the coding sequence of the *oculocutaneous albinism 2* (*oca2*) gene [11,12]. The finding that albinism is associated with shortened sleep raises the possibility that mutation of *oca2* also contributes to the evolution of sleep loss. To test this directly, we quantified sleep in surface fish with a CRISPR/Cas9 engineered deletion in exon 21 of the *oca2* gene (*oca2*^*Δ2bp*^). A deletion of this entire exon is found in Molino cavefish [11], suggesting the engineered mutation could phenocopy the naturally occurring mutation. This engineered mutation, when homozygous, results in complete loss of melanin pigmentation in surface fish, phenocopying albino cavefish (Fig 3A,B, [12]). Sleep was significantly reduced during the day and night in *oca2*^*Δ2bp/Δ2bp*^ surface fish compared to pigmented, wild-type *oca2*^*+/+*^ control siblings (Cohen’s d=1.3, a large effect, Fig 3,A-D). This reduction in sleep derives from reduced bout duration, which is significantly reduced in *oca2*^*Δ2bp/Δ2bp*^ surface fish compared to wildtype *oca2*^*+/+*^ control siblings (Fig 3E). Mutation of *oca2* did not result in a statistically significant change in bout number (Fig 3F). Together, these findings demonstrate that mutations in the *oca2* gene can affect both pigmentation and sleep, and raise the possibility that pleiotropy of the *oca2* gene contributed to the evolution of both albinism and reduced sleep in cavefish.

**Figure 3.**
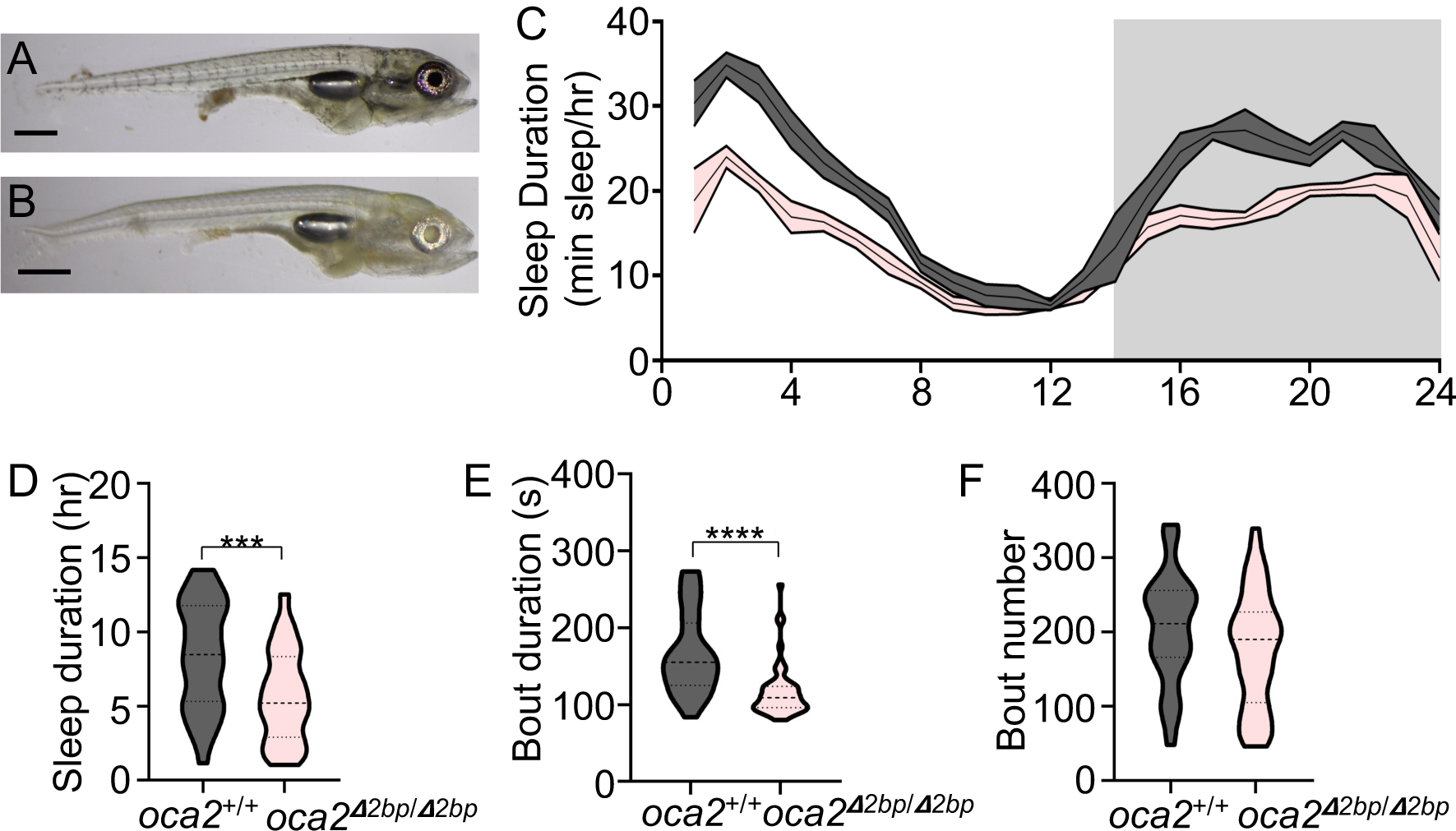
*oca2* mutant surface fish sleep less. (A) Pigmented wild-type and sibling (B) albino *oca2*^*Δ2bp/Δ2bp*^ 22 dpf surface fish. (C) 24-hour sleep profile of oca2^+/+^ and *oca2*^*Δ2bp/Δ2bp*^ surface fish siblings depicting the amount slept at 10-minute intervals over a 24-hour time frame. (D) Total sleep duration in *oca2*^*Δ2bp/Δ2bp*^ compared to wild-type oca2^+/+^ siblings (t-test, t=3.71, p=0.0004). (E) Bout duration in *oca2*^*Δ2bp/Δ2bp*^ compared to wild-type oca2^+/+^ siblings (Mann-Whitney, U=309, p<0.0001). (F) Bout number in *oca2*^*Δ2bp/Δ2bp*^ compared to wild-type oca2^+/+^ siblings (Mann-Whitney, U=662, p=0.092). For (C)-(F), pigmented wild-type (n = 32), and albino *oca2*^*Δ2bp/Δ2bp*^ (n = 53).

Surface fish heterozygous at the *oca2* locus (*oca2*^*Δ2bp/+*^) are pigmented and presented an intermediate total sleep duration phenotype which did not differ significantly from sleep in wild-type or *oca2* mutant siblings (Rerun with heterozygous individuals included: Average total sleep duration for *oca2*^*+/+*^ *=* 8.42 hours (n=32), *oca2*^*Δ2bp/+*^ = 7.03 hours (n=47), *oca2*^*Δ2bp/Δ2bp*^ = 5.64 hours (n=53); One-way ANOVA: F= 6.89, p= 0.0014; Tukey’s posthoc test: *oca2*^*+/+*^ vs *oca2*^*Δ2bp/+*^ (p=0.1674), *oca2*^*+/Δ2bp*^ vs *oca2*^*Δ2bp/Δ2bp*^ (p=0.1104), *oca2*^*Δ2bp/Δ2bp*^ vs *oca2*^*+/+*^ (p=0.0010)). To confirm that *oca2* regulates sleep in cavefish, we investigated whether our engineered loss-of-function *oca2* alleles in surface fish complemented naturally occurring deletions in *oca2* in albino cavefish. We crossed surface fish heterozygous at the *oca2* locus (*oca2*^*Δ2bp/+*^) to Pachón or Molino cavefish, both of which harbor natural coding mutations in *oca2* [11]. The presence of the engineered *oca2*^*Δ2bp*^ allele resulted in albino offspring in surface-Pachón and surface-Molino crosses (genotype *oca2*^*Δ2bp/ΔPa*^ and *oca2*^*Δ2bp/ΔMo*^, respectively), whereas offspring that inherited the wild-type allele from the surface parent (genotype *oca2*^*+/ΔPa*^ and *oca2*^*+/ΔMo*^) were pigmented (Fig 4A,B,D,E), confirming this mutation is sufficient to induce albinism in multiple cave populations and consistent with previous studies [12]. Further, albino *oca2*^*Δ2bp/ΔPa*^ and *oca2*^*Δ2bp/ΔMo*^ hybrid fish slept significantly less than their wildtype *oca2*^*+/ΔPa*^ and *oca2*^*+/ΔMo*^ hybrid pigmented siblings (Fig 4C,F). These findings demonstrate that the engineered *oca2* mutant allele fails to complement the sleep phenotype in Pachón and Molino cavefish, supporting the notion that loss of *oca2* contributes to sleep loss in multiple independently evolved cavefish populations. Bout duration was reduced in surface-Pachón *oca2*^*Δ2bp/ΔPA*^ hybrid fish compared to *oca2*^*+/ΔPA*^ siblings (Fig S3A). Both bout duration and bout number in surface-Molino *oca2*^*Δ2bp/ΔMo*^ hybrid fish were lower relative to siblings that inherited one wild-type surface allele, but they did not reach significance (Fig S3A,B,D,E). Finally, we crossed *oca2*^*Δ2bp/+*^ surface fish to Tinaja cavefish, which are not albino, and do not harbor known loss of function mutations in *oca2*, but which also present a reduced sleep phenotype [27]. We found no visible difference in melanin pigmentation between *oca2*^*Δ2bp/Ti*^ and *oca2*^*+/Ti*^ hybrid fish, confirming a lack of effect of mutations in *oca2* on the presence of melanin pigmentation in this population (Fig 4G,H). Further, sleep in the *oca2*^*Δ2bp/Ti*^ hybrid fish was not significantly different from sleep in the *oca2*^*+/Ti*^ siblings, suggesting *oca2* mutations do not play a role in sleep loss in this population (Fig 4I, Fig S3C,F).

**Figure 4.**
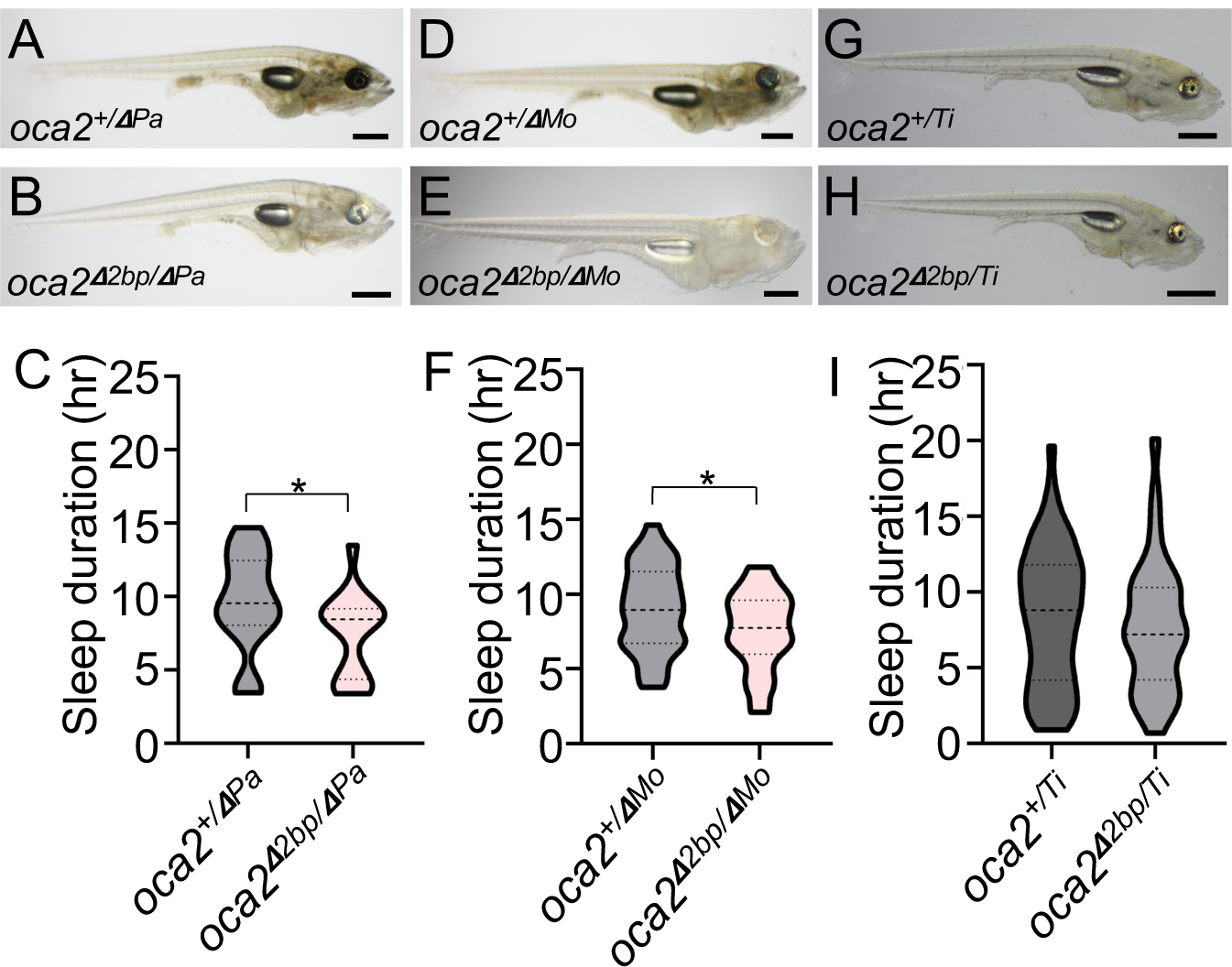
Lack of complementation in cave-surface F1 hybrid fish with two mutant *oca2* alleles. Melanin pigmentation (A, B, D, E, G, H) and sleep duration (C, F, I) were assessed in Pachon-surface F1 hybrid (A, B, C), Molino-surface F1 hybrid (D,E,F) and Tinaja-surface F1 hybrid fish (G,H,I). Images of 22 dpf (A) *oca2*^*+/ΔPA*^ (B) *oca2*^*Δ2bp/ΔPA*^ (D) *oca2*^*+/ΔMo*^ (E) *oca2*^*Δ2bp/ΔMo*^ (G) *oca2*^*+/Ti*^ (H) *oca2*^*Δ2bp/Ti*^ F1 hybrid fish. Total sleep duration in (C) *oca2*^*+/ΔPA*^ (n=21) compared to *oca2*^*Δ2bp/ΔPA*^ (n=17) siblings (t-test, t=2.210, p=0.033), (F) *oca2*^*+/ΔMo*^ (n=38) compared to *oca2*^*Δ2bp/ΔMo*^ (n=32) siblings (t-test, t=2.231, p=0.029) and (I) *oca2*^*+/Ti*^ (n=56) compared to *oca2*^*Δ2bp/Ti*^ (n=33) siblings (t-test, t=0.7128, p=0.478).

A central question in evolutionary biology is the extent to which selection or drift has driven trait evolution. Our findings that a single gene contributes to both albinism and sleep loss in multiple populations raises the possibility that selection for one or both of these traits leads to evolution of both traits in cave populations harboring *oca2* mutations. To test directly if *oca2* is under selection in cavefish, we conducted genome scans for selection using hapFLK (v1.4), software that allows for detection of genomic signatures of selection based on population genotyping [7]. The hapFLK statistic provides a powerful approach to detect regions of the genome under selection by testing for differentiation among populations in haplotype cluster frequencies exceeding what is expected by neutral evolution. We analyzed whole genome resequencing data from individuals of multiple populations of cave and surface fish collected previously [5]. In addition to cavefish populations from Molino (n=9), Pachón (n=10) and Tinaja (n=8), we analyzed surface *A. mexicanus* populations from two different localities: Rascón (n=7) and the Río Choy (n=9), the surface population used for behavioral data collection in this study. We also included a single individual from *Astyanax aeneus*, a closely related species, to serve as an outgroup. We observed statistically significant hapFLK values (1% FDR cutoff, p-values < 4.11e-05) within the *oca2* region, consistent with positive selection at this locus (Fig S4A). To identify the populations that experienced positive selection, branch lengths were re-estimated by building a local population tree for *oca2* using Reynolds distances based on haplotype frequencies. We observed significant p-values in branches corresponding to both surface and all three cave populations, Pachón, Molino, and Tinaja, (Fig S4B, Table S1), indicative of selection. This analysis reveals that population-specific *oca2* alleles are under positive selection across *A. mexicanus* populations. Therefore, these findings support the notion that selection for loss of *oca2* is a critical contributor to the evolution of sleep loss and albinism in cave habitats.

## Discussion

Understanding the relationships between evolved traits is a central goal of evolutionary biology. The interfertility of independently evolved *A. mexicanus* populations provides a system to investigate the genetic and functional relationships between traits. Previous studies of morphological trait evolution in *A. mexicanus* found that quantitative trait loci (QTL) for many seemingly unrelated morphological traits cluster in the genome more than expected by chance [22,28]. Further, evolved differences in sensory systems are thought to be critical drivers of the behavioral changes observed in cavefish, and in some cases, QTL for sensory traits and behaviors overlap [29,30]. However, none of these studies have addressed which genes and genetic lesions underlie these relationships. Thus, it is still unclear if these observed relationships are due to linkage, pleiotropy, or both. The finding that a single gene, *oca2*, underlies albinism and contributes to the reduction in sleep in two independently evolved cave populations of *A. mexicanus*, definitively demonstrates a pleiotropic role of a naturally occurring genetic variant in a morphological and a behavioral trait. To our knowledge, these findings represent the first association between monoallelic albinism and the evolution of an ecologically-relevant behavior in any cave animal.

The protein encoded by *oca2* functions at the first step of the melanin synthesis pathway in *A. mexicanus*, the conversion of L-tyrosine to L-DOPA [12,31]. Further, mutations in *oca2* have been linked to catecholamines. Morpholino knockdown of *oca2* in larval surface fish increases levels of dopamine, and albino cavefish have evolved higher levels of the catecholamines dopamine and norepinephrine [32–34]. This raises the possibility that OCA2 could modulate behaviors through regulating catecholamine levels [32,34]. Catecholamines regulate a number of behaviors, including sleep, feeding and social behaviors [35–37]. Further, many of these behaviors have evolved in cavefish [4,18,30,38] and reductions in schooling behavior, anesthesia resistance, and loss of sleep have all been linked to catecholamines in *A. mexicanus* through pharmacological analyses [30,34,39]. Thus, our observations that *oca2* contributes to sleep loss in cavefish could be due to effects of *oca2* on catecholamine levels. Further analyses to determine if loss of function mutations in *oca2* are sufficient to alter catecholamines are needed to examine this relationship.

Multiple adaptive roles for pigmentation have been shown across a variety species, including protection against ultraviolet radiation, parasitism, and thermoregulation [40]. In line with this, we found that *oca2* shows signatures of positive selection in both populations of surface fish. Further, all three cave populations examined here also show signatures of positive selection at the *oca2* locus. That a signature of selection is present in all populations regardless of whether they are albino or not suggests two non-mutually exclusive possibilities: First, that pigmentation is under selection across populations and is an adaptive trait in surface fish, offering protection against harmful UV radiation or thermal regulation. Second, other traits controlled by *oca2* are under selection across populations, supporting the hypothesis that *oca2* is pleiotropic.

While adaptive roles for pigmentation are well documented in surface populations, the evolutionary drivers of reductions of pigmentation in subterranean habitats are poorly understood. In cave populations, reduction or loss of melanin pigmentation in *A. mexicanus* cave populations has occurred through at least three mechanisms: reduction in the number of melanin producing melanophores, reduction in amount of melanin pigment produced, and the complete loss of melanin pigmentation, albinism, caused by mutations in *oca2* [11,16,17,41]. For decades, it has been argued that loss of pigmentation in cave animals is due to reduced selective pressure to maintain pigmentation within the dark cave environment [16,21]. Further, previous studies on the effects of QTL for melanophore number suggest that reductions of pigmentation alone may not be under direct selection [21]. Here, we provide genomic evidence that *oca2* is under selection in cave populations of *A. mexicanus*. Further, we demonstrate that loss of function alleles of *oca2* reduce sleep. Together, these results support the hypothesis that albinism has evolved through selection, not drift, in cave populations, and raise the possibility that selection has acted on another trait, distinct from albinism, sleep.

In conclusion, we report three complementary findings that suggest *oca2* contributes to sleep loss in cavefish. First, in F2 hybrid fish, sleep is lower in albino fish compared to pigmented fish from the same brood. Second, CRISPR-mediated mutation of *oca2* in surface fish reduces sleep. And finally, these CRISPR-mediated mutations fail to complement the sleep phenotypes of Pachón and Molino cavefish that harbor endogenous *oca2* mutations. Finally, we find that *oca2* is under selection in two albino cavefish populations, Pachón and Molino, as well as the hypopigmented Tinaja population. These findings raise the possibility that selection drove the evolution of albinism and sleep loss observed in multiple independently evolved cavefish populations.

## Materials and Methods

### Husbandry

Animal husbandry and breeding were carried out as described previously [42]. All procedures were done in accordance with the IACUC committee at Florida Atlantic University. Adult fish were housed in a fish facility with an Aquaneering flow-through system maintained at 23°C with a 14:10 hr light: dark cycle. Fish were bred by feeding frozen blood worms 2-3 times for the duration of breeding, and heating the water to 26-28°C to induce spawning. Larvae were raised in groups of 50 in stand-alone tanks, and fed brine shrimp once daily until 20 days post fertilization (dpf) for all assays described here.

### *oca2* mutant fish

The *oca2*^*Δ2bp*^ allele was previously isolated following CRISPR/Cas9 induced mutagenesis at the *oca2* locus [12]. Founder fish from this cross were from a surface fish population originated in Texas. These fish were outcrossed for 1-2 generations to surface fish from Mexico (Río Choy). For the experiments described here, we crossed heterozygous (*oca2*^*Δ2bp/+*^) F2 or F3 to one another, or crossed *oca2*^*Δ2bp/+*^ F2 or F3 to cavefish. All crosses were performed using pairs of fish, with each assay being performed on fish from multiple crosses.

Genotyping was performed as described previously [12,43,44]. DNA was isolated from whole larval fish or from adult fin clips by incubating larvae or fin clips in 50 mM NaOH for 30 minutes at 95°C. Following addition of 1/10^th^ volume Tris-HCl pH 8.0, PCRs were run using forward primers specific to the alleles: 5’-CTGGTCATGTGGGTCTCAGC-3’ was used for the wild type surface *oca2* allele and 5’-TCTGGTCATGTGGGTCTCATT-3’ was used for the 2 base pair mutant allele. The same reverse primer, 5’-TGTCAAGATATGTGATCTTTGGAAA-3’ was used for both reactions.

### Hybrid fish

To obtain F2 hybrid fish, a single surface fish female from a Mexican population was crossed with a single Pachón cavefish male to obtain cave-surface F1 hybrids. A single pair of F1 hybrid fish was crossed to produce all of the F2 hybrids described here. Wild-type surface and cave larvae were produced by group crosses of surface fish and Pachón cavefish. F1 fish for behavioral assays were progeny of a single pairwise cross between a female Río Choy surface fish and a male Pachón cavefish.

### Phenotyping

Sleep experiments were carried out as described previously [27,45]. Briefly, 20 dpf larvae were acclimated in 24-well plates for ∼15 hours prior to recording. Following feeding with artemia for 10 minutes, fish were recorded for 24 hours (14L:10D) starting at ZT0, lit from the bottom with LED white and IR lights. Videos were recorded at 15 frames per second using the video capturing software, VirtualDub (Version 1.10.4).

Videos were then subsequently tracked using Ethovision XT 13.0 (Noldus, IT) software. Sleep behavior parameters were defined from raw data using a custom-written Perl script and MS Excel macros. Sleep is defined by periods of inactivity during which individuals experience an increased arousal threshold [2]. Periods of inactivity that were of 60 seconds or greater were defined as sleep, as this period of immobility is associated with an increase in arousal threshold in *A. mexicanus* [4]. Total sleep duration, number of sleep bouts, and sleep bout length were quantified for each fish.

At 21 dpf, following the 24-hour sleep recording, fish were analyzed for feeding behavior. They were transferred to a 24-well plate pre-filled with approximately 70 *Artemia* naupili and were well and recorded for two hours. Total feeding was quantified by counting artemia before and after the feeding using Fiji [46] and subtracting to get the total number of artemia consumed.

Following feeding, fish were fasted for 20 hours (to limit autofluorescence), and at 22 dpf, fish were stained with Nile red (Sigma Aldrich 19123). The stock solution was prepared by dissolving Nile Red in acetone at a concentration of 1.25 mg/mL and stored in the dark at -20°C. Prior to staining the stock solution was diluted with fish water to a final working concentration of 1/1000. Fish were stained in a 24-well plate with 1mL of the working solution in each well, and placed in a 28°C incubator for 30 minutes, as previously described [19]. Following staining, fish were euthanized in 100 ug/mL MS-222 and imaged on a Nikon SMZ25 stereoscope using an GFP filter. Fish were scored for presence or absence of Nile Red staining.

Measurements of standard length and eye diameter were taken from these images, using Fiji [46]. Eye diameter was measured from anterior edge of the eye to the posterior edge. Standard length was measured from snout to caudal peduncle. Eye diameter throughout the paper was corrected for standard length by dividing eye diameter by standard length.

Color images of individuals of a subset of representative larval fish taken at 21 dpf using a Canon Rebel t6i camera on a clear background, illuminated from above and below.

### Statistical analysis for trait comparisons

All statistical analyses were performed using Graphpad Prism 8.4.3. All data was tested for normality using Shapiro-Wilk tests. Data which did not pass the normality test were analyzed using the nonparametric Kruskal-Wallis test. Where statistical significance was indicated, post-hoc comparisons were completed using Dunn’s multiple comparisons test. Normally distributed data with multiple groups were analyzed using an ANOVA, with post-hoc comparisons completed using a Tukey post-hoc test. For continuous traits which did not pass the normality test, relationships between these continuous traits were analyzed using the Spearman’s rank correlations. Relationships between binary and continuous traits were analyzed for normality, and then analyzed using a t-test if they passed the normality test, or a Mann-Whitney test if they did not pass the normality test. For adipose proportion, data error bars were calculated using z*-value of 1.96 and denote the margin of error of the sample proportion and data was analyzed using Fisher’s Exact Test, and a post-hoc of pairwise Fisher’s Exact Test was performed where significance was indicated. Effect size tests were also performed on F2 and *oca2* comparisons, an odds ratio effect size test was performed for adipose proportion data, all other effect size tests for comparisons were Cohen’s D-test.

### Test for selection

To test for signatures of selection on the *oca2* gene we used hapFLK (v1.4) [7] on *A. mexicanus* whole genome resequencing data from two surface populations, Rascón (n=7) and Río Choy (n=9), and three cave populations, Tinaja (n=8), Molino (n=9), and Pachón (n=10). We also included a single *Astyanax aeneus* surface fish as an outgroup. Details on sequencing and genotyping can be found in Herman et al. (2018) [5]. Briefly, samples were sequenced as 100 bp paired end reads on an Illumina HiSeq2000 at The University of Minnesota Genomics Center. Raw sequencing data for these samples was downloaded from NCBI (available under SRA Accession Numbers SRP046999, SRR4044501, and SRR4044502). Three samples (Rascon_6, Tinaja_6, and Tinaja_E) that were included in [5] were excluded from this analysis due to putative recent hybrid ancestry.

The haplotype-based hapFLK statistic is an extension of the SNP-based FLK statistic [7]. hapFLK provides a powerful approach to detect regions of the genome under selection by using a model that incorporates linkage disequilibrium to test for differentiation in haplotype clusters among populations. Unlike F_ST_, the hapFLK statistic accounts for hierarchical population structure and for the effects of recombination [7].

We first ran hapFLK in SNP mode to calculate FLK tests and to generate a kinship matrix based on the entire genome. The whole genome kinship matrix was then used to calculate haplotype-based hapFLK statistics on a 5 Mb region of chromosome 13 (35 Mb - 40 Mb) containing *oca2* (37,904,635-38,048,888 bp). For the haplotype-based tests, we used a K of 10 based on cross validation with fastPHASE (v1.4.8) and ran 20 expectation maximization iterations. hapFLK P-values were calculated using a chisquare distribution with the script scaling_chi2_hapflk.py. We used the R package qvalue to set a q-value cutoff of 0.01 and apply for a 1% false discovery rate (FDR) correction based on p-values present within the 35 Mb - 40 Mb regions of chromosome 13. To identify population-specific signatures of selection at *oca2*, we built local population trees using Reynolds distances based on haplotype frequencies with the script local_reynolds.py. We visualized changes in haplotype clusters across *oca2* in each population using the script hapflk-clusterplot.R. P-values were computed with the script local_trees.R by comparing the Reynolds distances among populations for the local tree compared to those from the whole genome tree. R and python scripts used in this analysis were downloaded from the hapFLK developers’ website https://forge-dga.jouy.inra.fr/projects/hapflk/documents.

## Acknowledgements

This work was supported by funding from NIH award 1R01GM127872-01 to SEM, ACK and NR and NSF awards DEB1754231 to JEK and AK, IOS165674 to ACK, and IOS1923372 to JEK, SEM, ED and NR and IOS1933076 to JEK, SEM and NR. All authors are grateful to Arthur Lopatto and Peter Lewis (FAU) for assistance with cavefish husbandry.

