## Supplemental documents for "Pleiotropic function of the *oca2* gene underlies the evolution of sleep loss and albinism in cavefish"

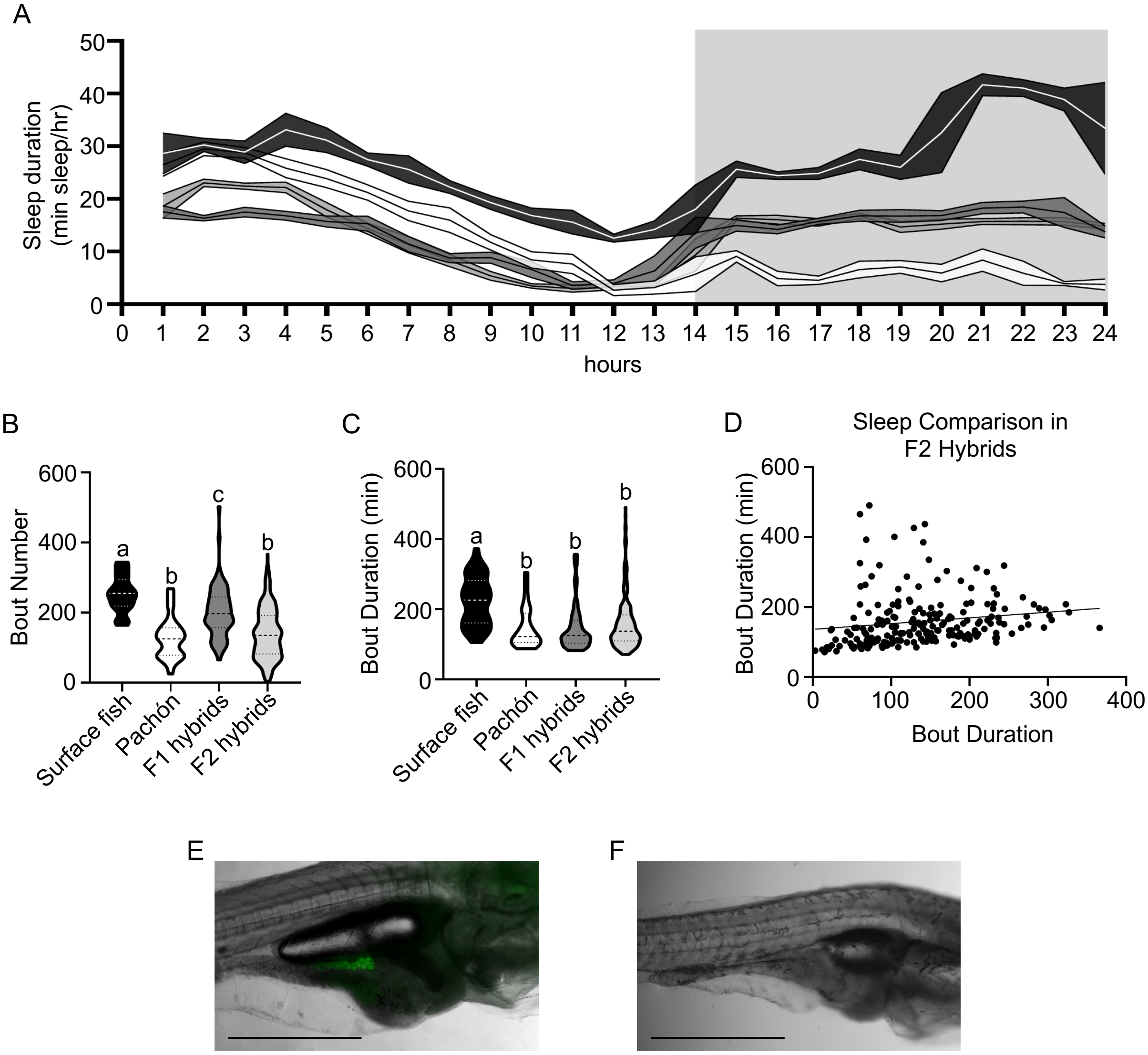


Figure S1 – Trait analysis in surface, Pachón, surface-Pachón F1 and surface-Pachón F2 fish.

(A) 24-hour sleep profile depicting the amount slept at 1-hour intervals over a 24-hour time frame. Black = surface fish, Grey = F1 hybrids, White = Pachón cavefish. Shaded areas of the graph represent when lights were off. (B) Bout number (Kruskal-Wallis: $H_2$=94.36, $p<0.0001$; Dunn's multiple comparisons post hoc test: SF vs. F1: z=3.243, p = 0.0071, Pa vs. F2: p>0.9999. All other: p<0.0001). (C) Bout duration (Kruskal-Wallis: $H_2$=37.35, $p<0.0001$; Dunn's multiple comparisons post hoc test: F1 vs. F2: z=1.418, p=0.9372, Pa vs. F1 and Pa vs. F2: p>0.9999. All other: p<0.0001). (D) Comparison of bout duration and about number in F2 hybrid crosses. (Spearman-rank correlation: r=0.3604, p<0.0001; linear regression is included as a descriptive: slope=0.1636.) (E) Image of 22dpf fish with adipose where Nile Red stain is pseudocolored green (scale=1mm)). (F) Image of 22dpf fish without adipose (scale=1mm). Graphs (B) and (C) are representations of median ± quartile.

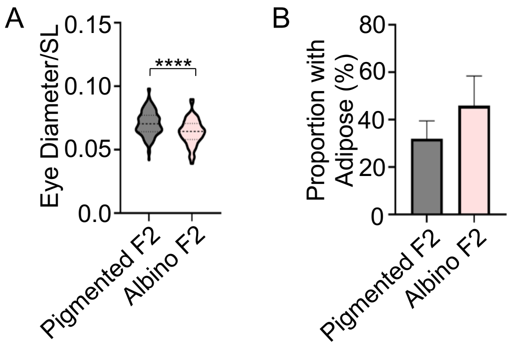


Figure S2 – Relationship between albinism and morphological traits in cave-surface F2 hybrid fish. (A) Eye diameter in albino vs. pigmented F2 hybrids, corrected for standard length (Mann-Whitney, u=6211, p<0.0001). This graph is a representation of median ± quartile. (B) Proportion of pigmented and albino F2 hybrid individuals with adipose. Fisher's Exact Test. Error bars calculated using z*-value of 1.96 and denote the margin of error of the sample proportion. (Fishers Exact tests: p=0.0805).

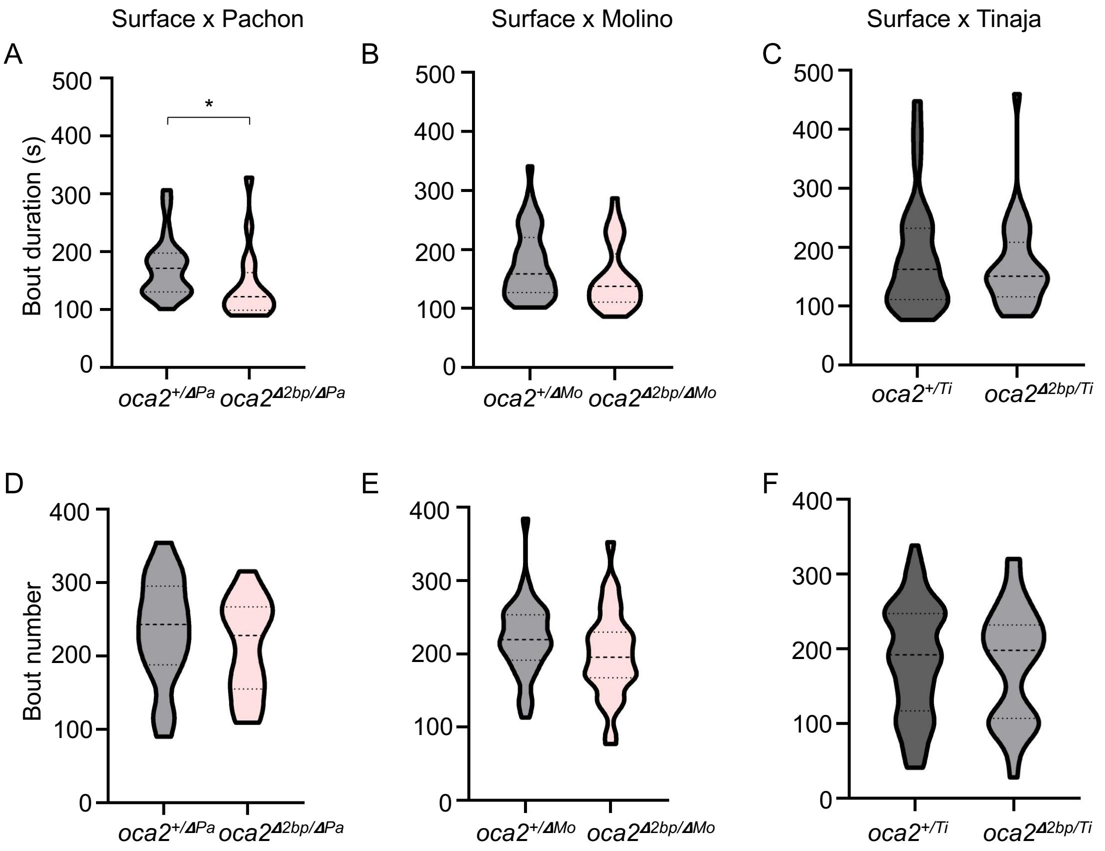


Figure S3 – Sleep architecture in surface-cave hybrid fish harboring an engineered *oca2* mutation. Average bout duration (A-C) and total number of bouts (D-F) were assessed in Pachón-surface F1 hybrids (A, D), Molino-surface F1 hybrid (B,E) and Tinaja-surface F1 hybrid fish (C,F). Average bout duration in (A) $oca2^{+/\Delta PA}$ (n=21) compared to $oca2^{\Delta 2bp/\Delta PA}$ (n=17) siblings (Mann-Whitney, U=93, p=0.0114). (B) $oca2^{+/\Delta Mo}$ (n=38) compared to $oca2^{\Delta 2bp/\Delta Mo}$ (n=32) siblings (Mann-Whitney, U=447, p=0.0581). (C) $oca2^{+/Ti}$ (n=56) compared to $oca2^{\Delta 2bp/Ti}$ (n=33) siblings (Mann-Whitney, U=884, p=0.7386). Number of bouts in (D) $oca2^{+/\Delta PA}$ (n=21) compared to $oca2^{\Delta 2bp/\Delta PA}$ (n=17) siblings (t-test, t=0.9381, p=0.3544). (E) $oca2^{+/\Delta Mo}$ (n=38) compared to $oca2^{\Delta 2bp/\Delta Mo}$ (n=32) siblings (t-test, t=1.631, p=0.1075). (F) $oca2^{+/Ti}$ (n=56) compared to $oca2^{\Delta 2bp/Ti}$ (n=33) siblings (t-test, t=0.2707, p=0.7873).

A

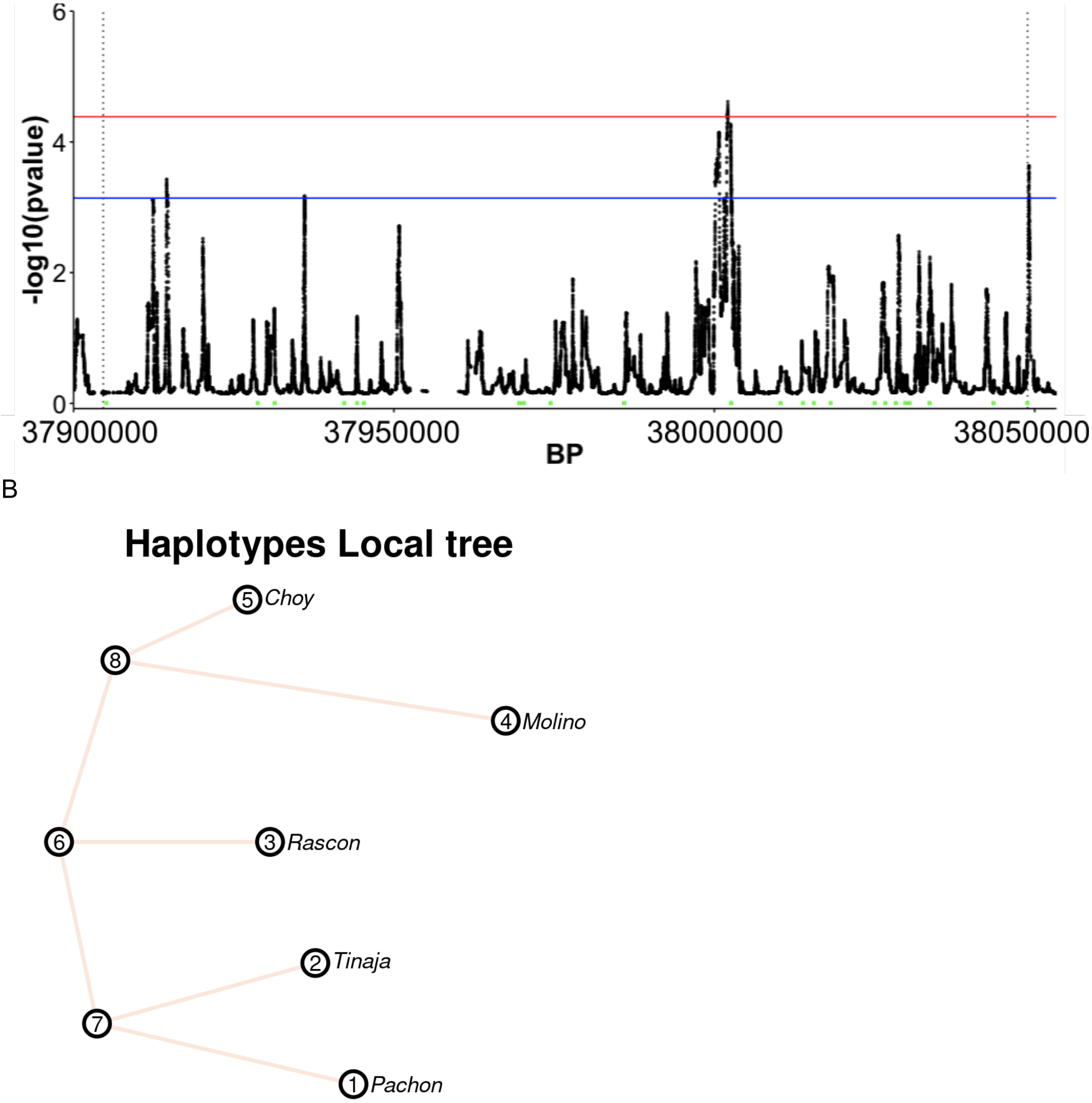


Figure S4 - Population genetics analysis at the *oca2* locus. (A) hapFLK p-values across *oca2* (region within dotted lines). Red line = 1% FDR cutoff. Blue line = 5% FDR cutoff. The 24 exons are shown in green boxes at the bottom of the plot. P-values were plotted along the antisense strand, so exon 24 is on the left end of the plot, near 37,900,000 bp, and exon 1 is on the right end of the plot near 38,050,000 bp. Peaks above the 1% FDR cutoff are present at exon 14 and a peak above the 5% FDR cutoff is present at exon 1.

(B) Local population tree for the *oca2* region of chromosome 13 (37,904,635-38,048,888 bp) using Reynolds distances based on haplotype frequencies.

Table S1 - P-values for branches within the haplotype-based population tree. Branch segments occur between nodes 1-8, corresponding to those shown in Supplementary Fig 4.

| Branch | Std. Error | t-value | Pr(>\|t\|) | P-value |
|---|---|---|---|---|
| 8<->4 | -0.388 | 0.01322 | -29.36 | 0.000087 |
| 8<->5 | -0.118 | 0.01322 | -8.96 | 0.002900 |
| 7<->1 | -0.312 | 0.01322 | -23.63 | 0.000170 |
| 7<->2 | -0.266 | 0.01322 | -20.09 | 0.000270 |
| 6<->7 | -0.009 | 0.01536 | -0.57 | 0.610000 |
| 6<->3 | -0.215 | 0.01145 | -18.8 | 0.000330 |
| 6<->8 | -0.05 | 0.01536 | -3.25 | 0.048000 |
